# 16p11.2 locus decelerates subpallial maturation and limits variability in human iPSC-derived ventral telencephalic organoids

**DOI:** 10.1101/2022.08.22.504797

**Authors:** Rana Fetit, Thomas Theil, Thomas Pratt, David J. Price

## Abstract

Inhibitory interneurons regulate the activity of cortical circuitry, and their dysfunction has been implicated in Autism Spectrum Disorder (ASD). 16p11.2 microdeletions are genetically linked to 1% of ASD. However, there have been few studies of the effects of this microdeletion on interneuron development. Using ventral telencephalic organoids derived from human induced pluripotent stem cells, we investigated the effect of this microdeletion on organoid size, progenitor proliferation and organisation into neural rosettes, ganglionic eminence (GE) marker expression at early developmental timepoints and expression of the neuronal marker, NEUN at later stages. Early deletion organoids exhibited significantly greater variations in size with concomitant increases in relative neural rosette area and the expression of the ventral telencephalic marker, COUPTFII, with significantly increased variability in these properties. Cell cycle analysis revealed a significant increase in total cell cycle length caused primarily by an elongated G1-phase, the duration of which also varied significantly more than normal. Late deletion organoids increased their expression of the neuronal marker NEUN. We propose that 16p11.2 microdeletions increase developmental variability and may contribute to ASD aetiology by lengthening the cell cycle of ventral progenitors, promoting premature differentiation into interneurons.

**Summary Statement:** Using 3D-region-specific organoids, we demonstrate that 16p11.2 deletion increases variability and prolongs the cell cycle of human subpallial progenitors by lengthening their G1 phase.

## Introduction

Autism Spectrum Disorder (ASD) is a complex, pervasive neurodevelopmental condition that is characterised by core symptoms which include difficulties in social cognition and communication, repetitive behaviours and hypersensitivities to external stimuli (Association, 2015, Varghese et al., 2017). The extent of the symptoms varies from patient to patient (London, 2014) and a wide range of comorbidities has been associated with ASD (Canitano, 2007, Hawks and Constantino, 2020, Lai et al., 2019). Evidence suggests that the underlying mechanisms leading to ASD manifestations are a result of early disruptions in the second trimester of foetal development (Wang et al., 2014), the same developmental period when inhibitory cortical interneurons are specified. Therefore, investigating GABAergic interneuron development is particularly relevant to ASD (Marín, 2012, Takano, 2015). Moreover, excitatory/inhibitory imbalance due to interneuron dysfunction has long been considered an important underlying cause of ASD (Rubenstein and Merzenich, 2003, Marín, 2012, Hussman, 2001, Uzunova et al., 2016). Human post-mortem studies provide sufficient evidence to implicate GABAergic and glutamatergic dysfunction in the aetiology of ASD (Fetit et al., 2021).

During development, cortical interneurons arise from the ganglionic eminences (GE) of the ventral telencephalon, which is divided into three proliferative zones, medial, caudal and lateral (MGE, CGE and LGE), distinguished by their expression of different molecular markers. NKX2.1 is highly expressed in the MGE, which generates the largest fraction of the cortical interneurons in both humans and rodents. A smaller proportion of interneurons arise from the CGE, which is marked by abundant COUPTFII expression. The LGE, on the other hand, only makes a minor contribution to cortical interneuron production (Hansen et al., 2013, Ma et al., 2012, Kelsom and Lu, 2013, Yang et al., 2021). The cortical interneurons produced in the MGE and CGE migrate tangentially through the LGE to integrate with excitatory projection neurons in the cortex (Whalley, 2013).

Although the aetiology of ASD is not yet fully understood, several genetic and environmental factors are known to play a role in its onset and development (De Felice et al., 2015, Varghese et al., 2017). Large genomic copy number variants (CNVs) account for approximately 10% of ASD cases (Ramaswami and Geschwind, 2018, Tuzun et al., 2005). 16p11.2 microdeletions, spanning around 600 kb and encompassing 47 genes (Marshall et al., 2008, Pinto et al., 2010), are associated with a variable spectrum of neurocognitive phenotypes. These include ASD, intellectual disability, morbid obesity, macrocephaly, or epilepsy at varying degrees of penetrance (Shinawi et al., 2010, Bochukova et al., 2010, Bijlsma et al., 2009, Fetit et al., 2020, Szelest et al., 2021). 16p11.2 microdeletions are also one of the most common genetic linkages to ASD (Fernandez et al., 2010, Weiss et al., 2008).

The underlying molecular mechanisms linking the 16p11.2 deletion to ASD remain largely unknown and have been studied largely in rodent models (Pucilowska et al., 2015, Blumenthal et al., 2014). A number of studies investigating the roles of individual genes within the 16p11.2 locus, such as *MAPK3* (Pucilowska et al., 2015, Mazzucchelli et al., 2002), *QPRT* (Feldblum et al., 1988, Haslinger et al., 2018), *KCTD13* (Golzio et al., 2012) and *TAOK2* (De Anda et al., 2012), suggested possible dysregulation of progenitor proliferation, neuronal migration and cortical lamination (Packer, 2016, Casanova, 2014). Murine models lacking the syntenic region on chromosome 7F3 recapitulate some ASD-like behaviours (Portmann et al., 2014, Pucilowska et al., 2015, Horev et al., 2011, Ouellette et al., 2020, Lu et al., 2019, Angelakos et al., 2017) and exhibit enhanced progenitor proliferation (Pucilowska et al., 2015) and basal ganglia abnormalities (Lu et al., 2018; Portmann et al., 2014). In addition, evidence from other studies suggests that it is likely that multiple genes within the region interact through shared pathways, contributing to the variable clinical phenotypes (Jensen and Girirajan, 2019, Pizzo et al., 2019).

The advancement of induced-Pluripotent Stem Cell (iPSC) and genome editing technologies has enabled the use of patient-derived tissue to study rare genetic mutations and model complex neurodevelopmental disorders. Cerebral organoids are 3D-cell aggregates derived from PSCs and contain many of the cell types found in embryonic brains, locally organized and behaving similarly to cells found in vivo (Lancaster et al., 2013). To date, only one published study has used iPSC-derived cortical organoids to investigate the effects of the 16p11.2 deletion on cortical development (Urresti et al., 2021). In addition to recapitulating the patient macrocephalic phenotype, 16p11.2 patient-derived cortical organoids exhibited an excess of neurons and depletion of neural progenitors (Urresti et al., 2021).

To date, there are no reports of using region-specific ventral organoids to specifically address the effects of this deletion on interneuron development. Here, we demonstrate that ventral organoids harbouring the deletion are more variable in size than normal. They exhibit significant relative increases in neural rosette area, COUPTFII expression at earlier timepoints, and prolonged cell cycle primarily due to lengthening G1. All these properties are significantly more variable in deletion organoids. Our results suggest increased variability and accelerated maturation of ventral deletion organoids, which may result in the premature differentiation of ventral progenitors into interneurons.

## Results

### Off-target genetic variation between cell lines was limited

We adapted the protocol described by (Sloan et al., 2018) to generate ventral telencephalic organoids from heterozygous 16p11.2 CRISPR/Cas9-deletion and isogenic control iPSC lines all derived from the same parent line, GM08330, which is referred to as GM8 in this study (Tai et al., 2016). No significant off-target CNVs were observed in previous studies of these CRISPR/Cas9-generated clones (Tai et al., 2016, Sundberg et al., 2021, Lim et al., 2022).

We used Illumina CytoSNP array analysis to confirm the presence of the 16p11.2 heterozygous deletion in all three deletion iPSC lines used in this study (**Fig. S1**). The parent line, GM8, and the three isogenic control lines did not carry any deletions or duplications of the 16p11.2 region. Our analysis also revealed a number of genomic locations where loss of heterozygosity (LOH) without a loss in copy number (copy neutral LOH) had occurred, usually in all lines (green bands, **Fig. S1**; **Table S2**. Of the few additional deletions and duplications that were present in some lines and not others (red, yellow, blue and purple bands, **Fig. S1; Table S2**), only five of the affected regions contained protein-coding genes, as follows. (1) A heterozygous deletion involved one protein-coding gene on chromosome 4 in control lines FACS52 and FACS53. (2) A heterozygous duplication involved six protein coding genes on chromosome 4 in deletion line DELD5. (3) The control line FACS53 had three additional CNVs: a heterozygous deletion on chromosome 5 involving one gene, a heterozygous duplication involving 3 genes on chromosome 4 and a heterozygous duplication involving 8 genes on chromosome 9. We designed our experiments to take into account the possibility that the effect of 16p11.2 deletion might be modified by these few heterozygous variants.

This study was divided into two parts. The first part investigated initial organoid growth between 5-25 days. The second part focused on the cell cycle kinetics of ventral progenitors and the differentiation of their daughter cells between days 33-70 **(Fig. 1A, Table S1).** Organoids grown for 33-35 days in Parts 1 and 2 were from three control lines (FACS51, FACS52 and FACS53) and two deletion lines (DELD5 and DELA3). Each of four separate batches of organoids (RF1-4) included at least one control and one deletion line. Across the four batches, all lines were replicated twice, except FACS53, which was replicated 3 times, and DELD5, which was replicated four times. All replicates included 10-15 organoids. For some experiments in Part 2, another batch of ventral organoids from the parent line (GM8) and the deletion line (DELB8) containing 4-6 organoids per line was maintained for 46-70 days (**Table S1)**. A combination of Linear Mixed Effects (LME) analysis, Welch’s ANOVA and Student’s t-test was used to account for the variability due to both batch-effects and cell line differences, and assess the statistical significance of our findings.

**Fig. 1:**
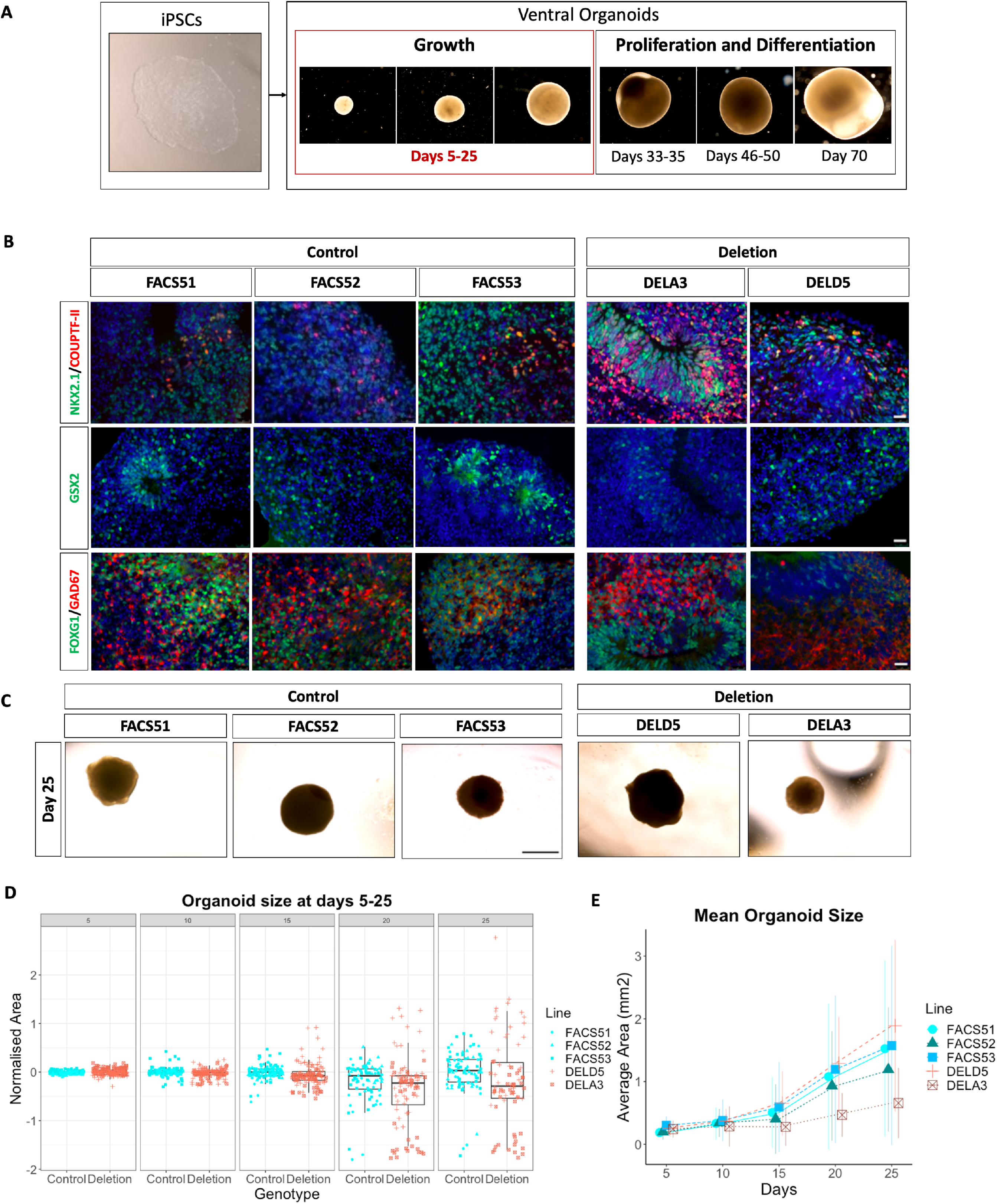
Ventral organoids express ventral telencephalic markers and exhibit variations in size. **(A)** Overview of the study design with representative images of organoids. The first part focused on organoid growth by measuring organoid size between 5-25 days. The second part focused on the proliferation of ventral progenitors and their differentiation by analysing cell cycle kinetics at 33-35 days and examining the expression of neuronal marker at 46, 50 and 70 days. (B) Both control and deletion organoids express the GE markers NKX2.1, COUPTFII and GSX2 as well as the forebrain marker FOXG1 and the GABAergic marker GAD67 at day 33-35. Images are shown merged with DAPI (blue), scale bar = 25 μm. (C) Representative images of the organoids at day 25. Scale bar = 1mm. (D) Projected area of ventral organoids over a period of 25 days. Box plots showing organoid area (mm^2^) normalised to the average of the control area for every batch at different timepoints. Data is pooled from the different replicates across four different batches. Every dot represents an individual organoid. Different shapes represent the different cell lines used. (E) Average organoid area (absolute values) for every cell line at the different timepoints. Error bars = (mean±2*s.d.). Sample size by genotype: day 5, control=152, deletion =149; day 10, control=137, deletion =131; day 15, control=114, deletion =115; day 20, control=83, deletion =86; day 25, control=85, deletion=74.

### Deletion and control organoids developed ventral telencephalic identity

We first characterised the organoids by immunohistochemistry (IHC) to assess their expression of forebrain and ventral telencephalic markers. At days 33-35, ventral organoids from both control and deletion lines expressed the GE markers NKX2.1, COUPTFII and GSX2, as well as the forebrain marker, FOXG1, and the GABAergic marker, GAD67 **(Fig. 1B).** These findings confirmed that the protocol used here indeed generated control and deletion organoids of ventral telencephalic identity.

### Early 16p11.2 deletion organoids exhibited abnormal variations in size

We assessed organoid growth because macrocephaly is frequently reported in 16p11.2 deletion patients (Qureshi et al., 2014) and increases in the volume of subcortical GE-derived brain structures, such as the striatum and globus pallidus, have been shown in 16p11.2 deletion mice (Rein and Yan, 2020). The projected area of ventral organoids from deletion lines (DELD5 and DELA3) and control lines (FACS51, FACS52 and FACS53) was measured every 5 days over a period of 25 days. **Fig. 1C** shows representative organoids from the different lines at day 25. **Fig. 1D** shows data on individual organoid areas normalised to the average area of the control organoids in the same batch at the same age **(Table S3).** The mean organoid areas for each line are shown in **Fig. 1E.** The organoids from the control lines all grew at relatively similar rates compared to the organoids from the deletion lines, whose sizes became more variable from day 15 on (F-test for comparing variance between genotypes, p= 2.30e-07, 4.12e-07 and 5.23e-10 for days 15, 20 and 25 respectively). This variation was between batches of deletion organoids rather than between deletion lines (**Fig.S2A**). Taking batch variability into account, LME analysis revealed no significant effects of the genotype on organoid area at day 25 (Type-III ANOVA, p=0.113). Our findings showed that 16p11.2 ventral organoids were not consistently larger or smaller but exhibited significantly greater variation in growth rates compared to their batch-matched controls.

### 16p11.2 deletion organoids exhibited increased potential for neural rosette formation

Given that several genes within the 16p11.2 deletion region are expressed in neural progenitor cells (NPCs) (Morson et al., 2021), we investigated whether the deletion affects the size and abundance of neural rosettes, which are radial arrangements of NPCs that resemble the developing neuroepithelium (Deglincerti et al., 2016, Townshend et al., 2020). We delineated the rosette structure based on the radial, circular arrangement of NPCs using DAPI (**Fig. 2A**). Neural rosettes often appeared larger and more abundant in organoids from both deletion lines compared to controls (**Fig. 2A**). We quantified the area occupied by the neural rosettes relative to the area of the organoid **(Table S5)**, thereby taking variations in organoid size into account. Compared to controls, significantly larger average relative rosette area was found in deletion organoids (Welch’s ANOVA, p= 0.000564; large effect size, Cliff’s delta = 0.6518519; **Fig. 2C**). Post-hoc comparisons of the individual lines are summarised in **Table S4.** Deletion organoids exhibited significantly greater variation in relative rosette area compared to their isogenic controls, with some deletion organoids in some batches generating very few rosettes (F-test, p= 5.382e-10; **Fig S2B**). There was no obvious relationship between whether deletion organoids increased their relative rosette area and their size (**Fig. S2C,** Spearman correlation R=−0.12, p=0.71). Collectively, our findings suggest that 16p11.2 deletion increased the potential of ventral organoids to form neural rosettes and increased the variation in rosette generation.

**Fig. 2:**
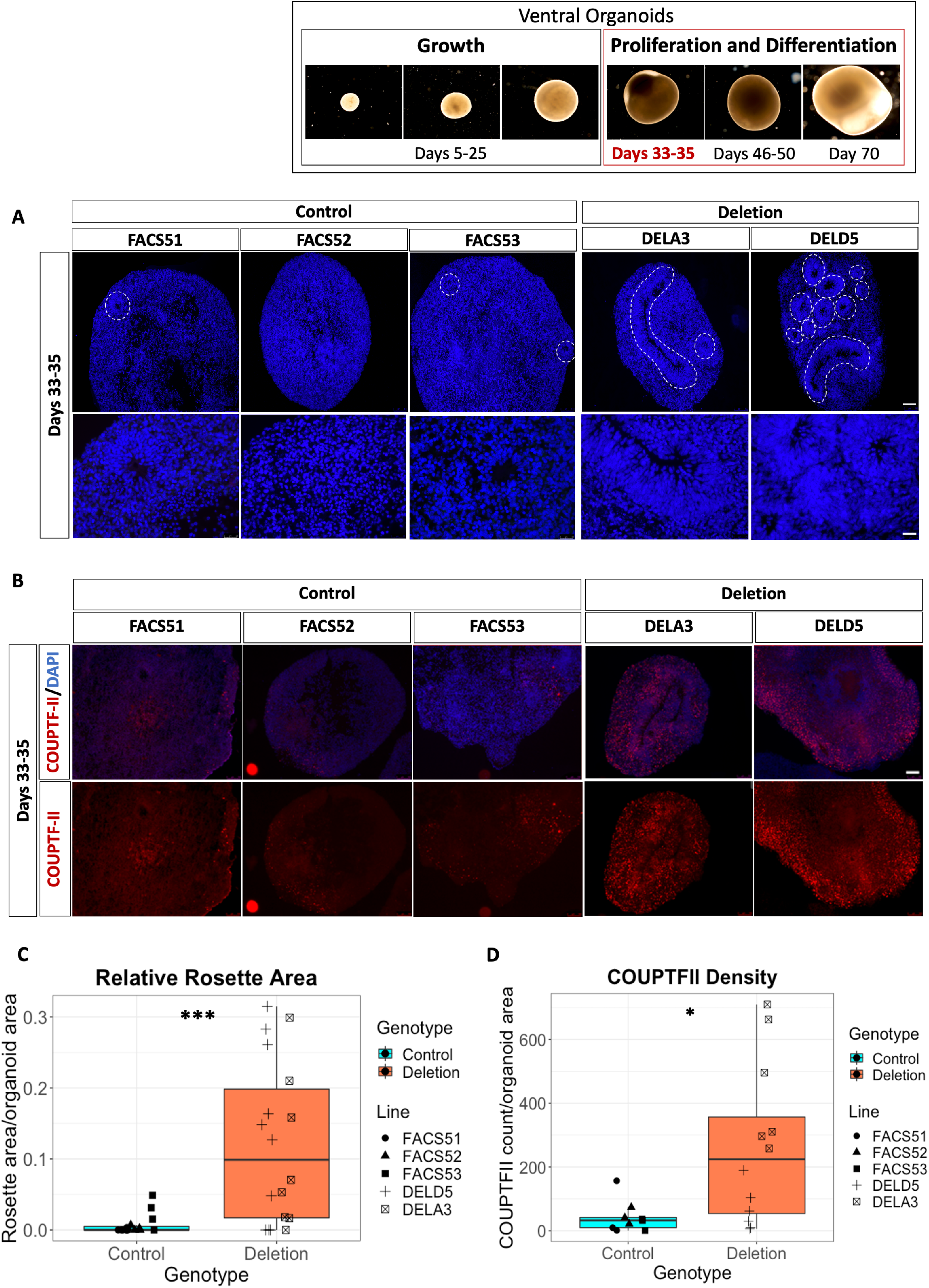
Quantification of neural rosettes and COUPTFII expression in ventral organoids at day 35. (A) Rosettes in representative ventral organoids from control and deletion lines outlined with dashed lines. Scale bar = 100μm (B) COUPTFII expression in control and deletion lines. Top panels show COUPTFII and DAPI, bottom panels show the red channel for COUPTFII only. Scale bar = 100μm (C) Boxplots showing the rosette area relative to organoid area grouped by genotype (control n=15, deletion n=18). Statistical significance determined using Welch’s ANOVA, p=0.000564. (D) Boxplots showing COUPTFII density (COUPTFII counts / organoid size). Data is grouped by genotype (control n=9, deletion=12). Statistical significance determined using Welch’s ANOVA, p=0.01093.

### 16p11.2 deletion organoids exhibited a significant increase in COUPTFII-expressing cell density

While characterising ventral organoids as described above, we noticed that COUPTFII was expressed by many more cells in organoids from both deletion lines than in controls **(Fig. 2B)**. The number of cells expressing COUPTFII was quantified and COUPTFII+ cell density was calculated, thereby taking into account the variations in organoid size. A significant increase in COUPTFII+ cell density was observed in the deletion organoids compared to their isogenic controls (Welch’s ANOVA, p= 0.01093; large effect size, Cliff’s delta= 0.5925926, **Fig. 2D).** Post-hoc pairwise-comparisons of the individual lines are summarised in **Table S4**. Deletion organoids exhibited significantly greater variation in COUPTFII+ cell density, with some deletion organoids in some batches generating relatively few COUPTFII+ cells (F-test, p= 9.968e-05; **Fig. S2C,D**). There was no relationship between whether deletion organoids increased their relative rosette area and their density of COUPTFII+ cells (**Fig. S2E)**. These findings suggest that ventral organoids harbouring 16p11.2 deletion have the potential to generate proportionately more COUPTFII+ cells and exhibit increased variability in COUPTFII+ cell density.

### 16p11.2 deletion did not alter the proportions of proliferating progenitors or immature neurons at early timepoints

Neural rosettes can respond to different patterning cues and initiate differentiation into region-specific neuronal fates (Elkabetz et al., 2008). Because more developed neural rosettes were observed in the deletion organoids, we then asked whether the deletion affected the proportions of proliferating progenitors and neurons in the organoids. We observed that SOX2, a marker associated with progenitor and relatively undifferentiated precursor cells (Pagin et al., 2021), was expressed in neural rosettes (white arrows, **Fig. 3A, B, D**) and cells surrounding them in the outer region of the ventral organoids (yellow arrows, **Fig. 3A, B**) and TUJ1 (green arrows, **Fig. 3C, D**), a marker expressed in immature, newly generated postmitotic neurons (von Bohlen und Halbach, 2007, Liu et al., 2007), was expressed by cells surrounding the rosettes in the outer regions of the organoids (yellow arrowhead, **Fig. 3C, D**).

**Fig. 3:**
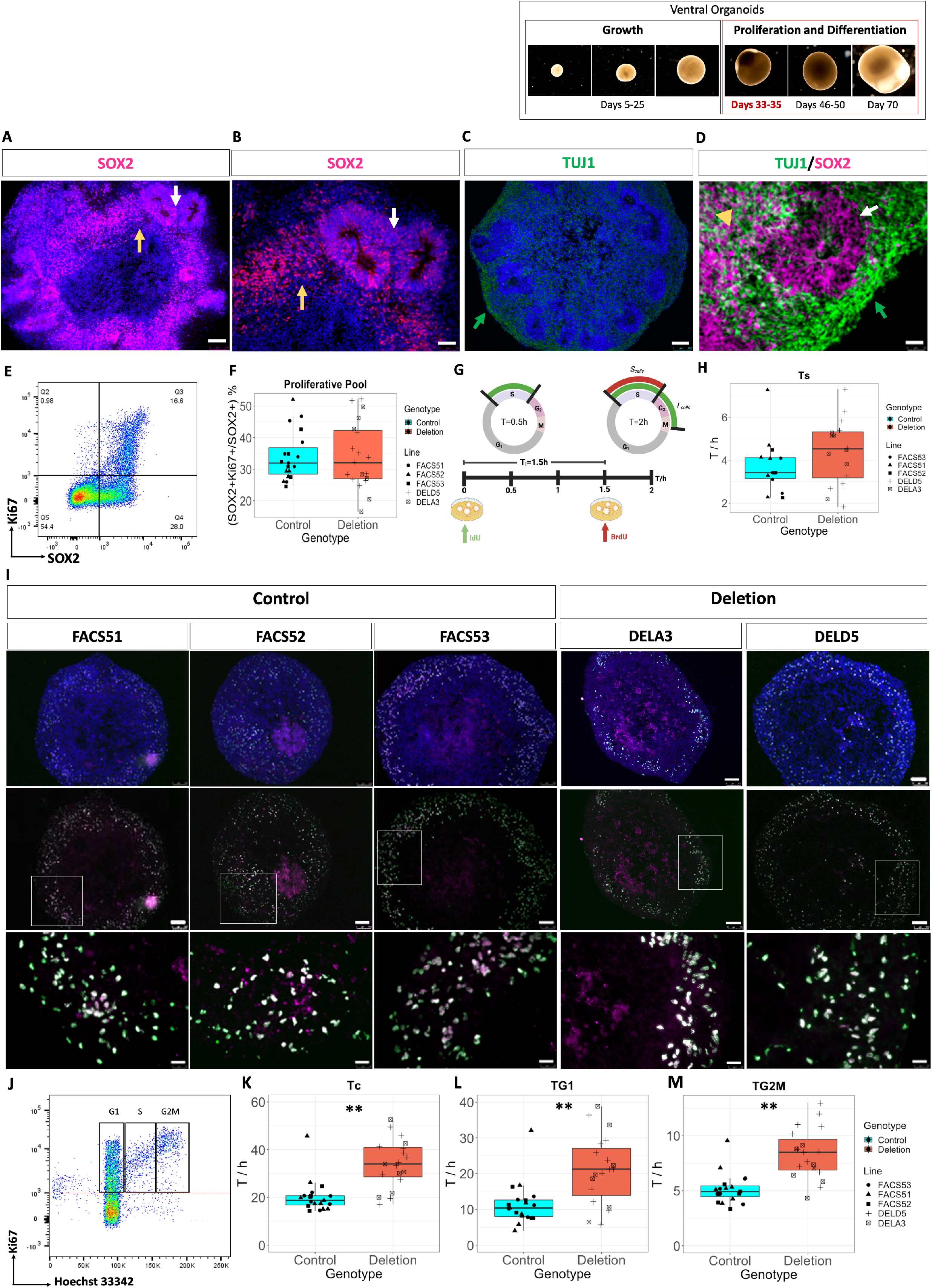
Cell cycle kinetics in ventral organoids. (A, B) SOX2+ cells (magenta) reside in the outer rim of the ventral organoids (yellow arrows), and in neural rosettes (white arrows). Scale bars = 100 μm (A) and 50μm (B). (C) Young, immature TUJ1+ neurons (green) reside outside neural rosettes in the organoid periphery (green arrow). (D) A neural rosette positive for SOX2 (magenta, white arrow) and surrounded by TUJ1+ neurons (green arrow). Some TUJ1+ cells also express SOX2 (yellow arrowhead). Scale bar = 25μm. (E) Flow cytometric density plot showing SOX2 (x-axis) against Ki67 (y-axis) in a representative organoid. In this organoid, 37% of the SOX2+cells are cycling and positive for the cell cycle marker Ki67. (F) Proportion of SOX2+Ki67+ progenitors as percentages of all SOX2+ cells in the organoid (proliferative pool). Sample size by genotype: n= 21 organoids for control and deletion. Sample size by cell line: FACS51 n=9, FACS52 n=5, FACS53 n=7, DELD5 n=14, DELA3 n=7. (G) Schematic representation of the IdU/BrdU double labelling experiment (created using BioRender.com). (H) Duration of S-phase (Ts) in hours (h) calculated from double IdU/BrdU labelling. (I) Representative images of ventral organoids stained with monoclonal antibodies specific for both both IdU and BrdU (green) and BrdU only (magenta) to allow for the identification of S_cells_ and L_cells_. Top panels show organoids at 10x merged with DAPI (blue). Middle panels show the green and magenta channels. Scale bars = 100 μm. White boxes outline the regions magnified in the lower panels. Scale bars = 25 μm. (J) Density plot of Ki67 (y-axis) against Hoechst 33342 (x-axis). SOX2+ cells in the cell cycle are those positive for Ki67 (above the dotted red line). Cells in G_1_ are those with 2n DNA content and those in G_2_M are those with double the DNA content, 4n. In between are cells in S phase. (K-M) Duration of the total cell cycle (Tc) and of the individual cell cycle phases in ventral organoids grouped by genotype in hours (h). Statistical significance determined using LME analysis to account for batch variability. Type-III ANOVA, p= 0.001733 for Tc, 0.008776 for TG1 and 0.001826 for TG2M. Sample size by genotype: control n=19, deletion n=19. Sample size by cell line: FACS51 n=8, FACS52 n=5, FACS53 n=6, DELD5 n=12, DELA3 n=7.

To determine proportions of proliferating progenitors and neurons, days 33-35 organoids were dissociated, fixed and permeabilised, then analysed with flow cytometry. Cells were labelled with TUJ1, SOX2 and the cell cycle marker Ki67 to unambiguously identify the cycling progenitor cells, which we will refer to as the proliferative pool **(Fig. S6)**. Not all SOX2+ cells expressed Ki67 (**Fig. 3E**) indicating the SOX2 expression persists after cells exit the cell-cycle, which was also observed for day 30 ventral organoids in a recently published study (Xiang et al., 2017) (**Fig. S3A**).

We first quantified the proportion of the SOX2+ cells that were proliferative (SOX2+Ki67+cells/SOX2+ cells) in our ventral organoids (**Fig. 3F, Fig. S3B**). We found no significant differences between deletion and control organoids, neither in average proportions (taking batch variability into account with LME analysis; Type-III ANOVA, p= 0.9748) nor in variability (F-test, p= 0.1977; **Fig. S3B**). We then quantified the proportion of TUJ1+ neurons and found no significant differences between deletion and control organoids **(Fig. S3C and D,** LME analysis; Type-III ANOVA, p= 0.3516). Moreover, no significant differences in the ratio of TUJ1+ neurons to SOX2+Ki67+ progenitors were found (**Fig. S3E and F**; LME analysis, Type-III ANOVA, p= 0.177). Our findings suggest that at this developmental timepoint (day 33-35), 16p11.2 microdeletion increases the potential of ventral progenitors to organise into neural rosettes without affecting the proportions of proliferative cells and immature neurons in the deletion organoids.

### 16p11.2 deletion organoids exhibited increased cell cycle lengths and lengthened G1 and G2M phases

Studies in 16p11.2 mouse models and patient derived NPCs have demonstrated enhanced cortical progenitor proliferation (Pucilowska et al., 2015, (Connacher et al., 2022), whereas other studies utilising human iPSC-derived cortical NPCs showed no difference in NPC proliferation rates (Deshpande et al., 2017, Roth et al., 2020). Therefore, we investigated whether the deletion affects the cell cycle of ventral progenitors within the outer SOX2+ peripheral region of the organoids, since control organoids formed very few rosettes. First, we determined the length of S phase (Ts) using double iododeoxyuridine (IdU) and bromodeoxyuridine (BrdU) labelling, as previously described (Martynoga et al., 2005) (**Fig. 3G,I, Table S5**). Although deletion organoids revealed a slight increase in the duration of S-phase (Ts) **(Fig. 3H),** this was not statistically significant (LME analysis, Type-III ANOVA, p= 0.1017). There was no significant difference in variation between control and deletion organoids (**Fig. S4A**, F-test, p= 0.517).

The numbers of SOX2+ cells that were proliferative, as marked by Ki67 expression, were calculated from flow cytometry (**Fig. 3E**), together with the number of these cells in the different cell cycle phases (**Fig. 3J**; **Table S6).** These data were used to calculate he total cell cycle length (Tc) and the duration of G1 (T_G1_) and G2M (T_G2M_) phases as described previously (Martynoga et al., 2005). The average total cell cycle length (T_C_) for the deletion organoids was 33.8 hours, which was significantly higher than that of the control organoids of 20.1 hours (**Table S6**; LME analysis, Type-III ANOVA, p= 0.001733; large effect size, omega_squared = 0.40, **Fig. 3K, Fig. S4B**). Moreover, significant increases in the duration of G1 (T_G1_) and G2M (T_G2M_) phases were observed in the deletion organoids, compared to controls (**Figs. 3L and M**, LME analysis, Type-III ANOVA, p= 0.008776 and 0.001826 for T_G1_ and T_G2M_, respectively). The effect sizes were large in both cases (Omega_squared = 0.37 and 0.83 for T_G1_ and T_G2M_ respectively). In addition, deletion organoids exhibited greater variations in T_G1_ and T_G2M_ compared to controls (**Fig. S4C and D**; F-test, p= 0.05052 and 0.04201 for T_G1_ and T_G2M_, respectively). These findings suggest an elongation of the cell cycle due to increased G1 and G2M phase length in deletion organoids concomitant with increased variability.

### Longer cell cycles correlated with lengthened G1 phase and increased relative rosette area in deletion organoids

We then asked whether the increased relative rosette area is correlated with longer Tc. Therefore, we calculated Tc for the organoids analysed in the imaging dataset and performed a correlation analysis between relative rosette area and Tc (**Table S5, Fig. 4A**). Organoids from the two deletion lines behaved similarly, in that those exhibiting high relative rosette area also exhibited longer Tc. Correlation analysis revealed a significantly stronger positive correlation between relative rosette area and Tc in deletion organoids than in controls, where a negative but insignificant correlation was observed (**Fig. 4A**, Spearman correlation, R=0.59 p=0.01 and R=−0.28 p=0.31 for deletion and controls, respectively). There was no evidence of a correlation between COUPTFII+ cell density and Tc (**Fig. S4E**).

**Fig. 4:**
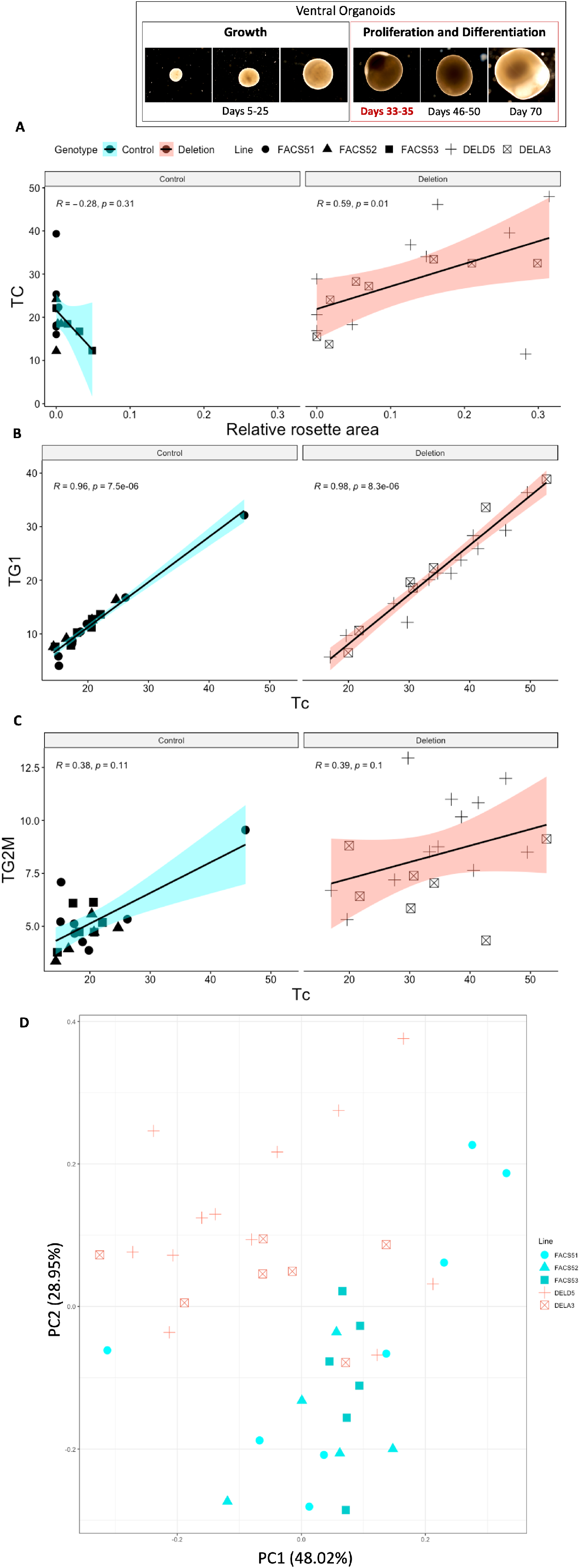
Linking relative rosette area, TG1 and TG2M to TC in imaging and flow cytometry datasets. (A) Correlation analysis between TC and relative rosette area in the imaging dataset (Spearman correlation). (B, C) Correlation analysis between TC and TG1 and TG2M respectively in the flow cytometric dataset (Spearman correlation). Control n=19, Deletion n=19. (D) PCA analysis of the organoids in flow cytometric dataset, grouped by cell line. Every individual point represents an organoid. Different shapes correspond to the different cell lines used.

After that, we examined the relationship between Tc, T_G1_ and T_G2M_ in the organoids analysed by flow cytometry (**Table S6**). We can clearly see that T_G1_ increases with longer Tc, a correlation that was both strong and significant in the deletion and control organoids (**Fig. 4B**, Spearman correlation, R=0.98 p=8.3e-06 and R=0.96 p=7.5e-06 for deletion and control organoids respectively). The correlation between T_G2M_ and Tc was rather moderate and insignificant (**Fig. 4C**; Spearman correlation, R= 0.38 p=0.1 and R=0.39 p=0.1 for control and deletion respectively).

We then ran a principal component analysis (PCA) using all variables generated in the flow cytometric dataset to observe how similar or different the individual organoids are within the control and deletion populations. PCA clearly separates most deletion organoids from the controls (**Fig 4D**). Taken together, we can conclude that the increase in relative rosette area in the deletion organoids correlates with longer Tc, which is primarily due to the lengthening of G1 phase. Moreover, the effects of the 16p11.2 CNV are similar in organoids from both deletion lines. **Tables S5** and **S6** list all the variables measured and calculated for every organoid in the imaging and flow cytometric datasets.

### 16p11.2 deletion increased NEUN expression at later stages

Because cell cycle and G1-phase lengthening are features of neural stem cells undergoing neurogenic cell divisions (Wilcock et al., 2007, Molina et al., 2020), we investigated whether this may affect progenitor differentiation in deletion organoids. We performed an additional study to generate one batch of ventral organoids from the parent line (GM8) and the deletion line (DELB8) containing 4-6 organoids per line. Organoids were maintained for longer than 35 days and the expression of NEUN, a marker of neuronal maturation that labels mature neurons (Gusel’Nikova and Korzhevskiy, 2015) was examined at days 46, 50 and 70. When we sampled from these organoid batches at day 35, they exhibited more neural rosettes (**Fig. S5A**), higher COUPTFII expression (**Fig. S5B**) and were positive for ventral telencephalic markers (**Fig. S5C**), thus recapitulating our findings in organoids from the deletion lines examined earlier.

A significant increase in *NEUN* mRNA expression was observed at day 46 in deletion organoids compared to their isogenic controls (**Fig. 5A**, p < 0.05; Student’s t-test). We then examined NEUN expression at day 50 by IHC to find a significant increase in the mean NEUN fluorescent intensity relative to organoid size in the deletion organoids (**Fig. 5B, C** and **Table S7**, p=0.01716; Student’s t-test). In contrast, no differences in *NEUN* mRNA expression were found at day 70 (**Fig. 5D**). Taken together, our findings suggest that the 16p11.2 deletion accelerates neuronal output in ventral organoids at later timepoints.

**Fig. 5:**
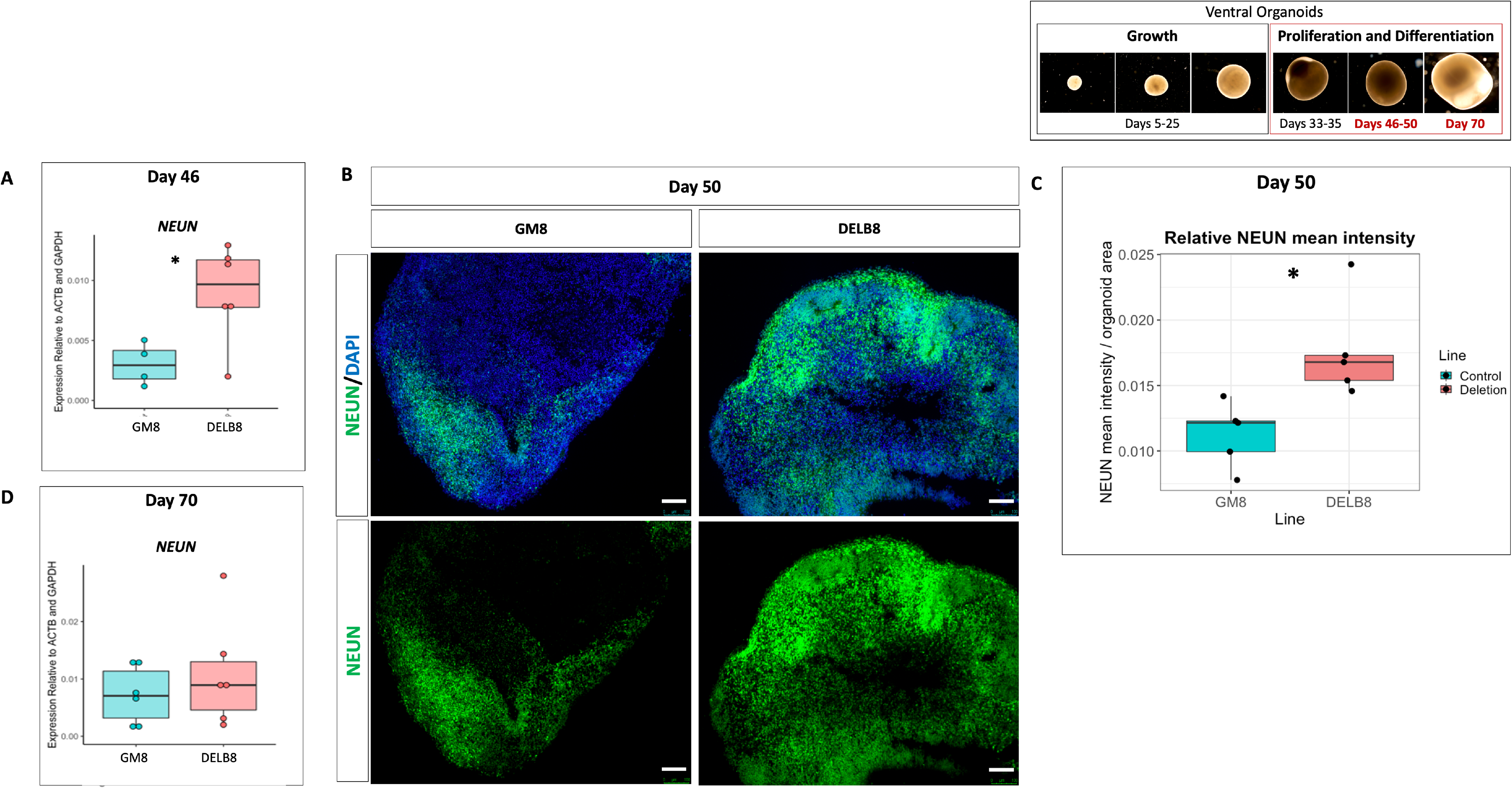
RT-qPCR and IHC analysis of NEUN expression in ventral organoids from the cell lines GM8 and DELB8. (A) Boxplots showing the relative mRNA expression of *NEUN* in control (green) and deletion (red) organoids at 46 days of differentiation. (B) NEUN expression at day 50 in representative control and deletion organoids. Top panels show NEUN and DAPI, bottom panels show cells expressing NEUN only. Scale bar = 100μm. (C) Mean NEUN intensity relative to organoid size at day 50. (D) Relative mRNA expression of NEUN at 46 days. N=4-6 organoids, * p < 0.05; Student’s t-test.

## Discussion

### Summary of the effects of 16p11.2 deletion in ventral organoids

To our knowledge, this is the first study to use ventral telencephalic organoids to investigate the effects of 16p11.2 deletion on early aspects of interneuron development in the ventral telencephalon. The first signs of an effect came from the increased variability in the growth of deletion organoids over the first few weeks of their development, although their average growth rates did not differ significantly from those of controls. Subsequently, at stages when telencephalic organoids start to organise their neural cells into rosettes in a process that resembles the development of the neuroepithelium of the neural tube, deletion organoids showed, on average, increased rosette formation. This increased state of organisation coincided with higher densities of COUPTFII expressing cells and longer cell cycles (Wilcock et al., 2007, Molina et al., 2020). As with the growth rates recorded at earlier stages, the relative rosette area, COUPTFII density and the duration of G1 and G2M phases varied more in deletion organoids than in controls. Our study also suggested that, at later stages, the deletion accelerated ventral neuronal production. These findings are summarised in **Fig. 6**.

**Fig. 6:**
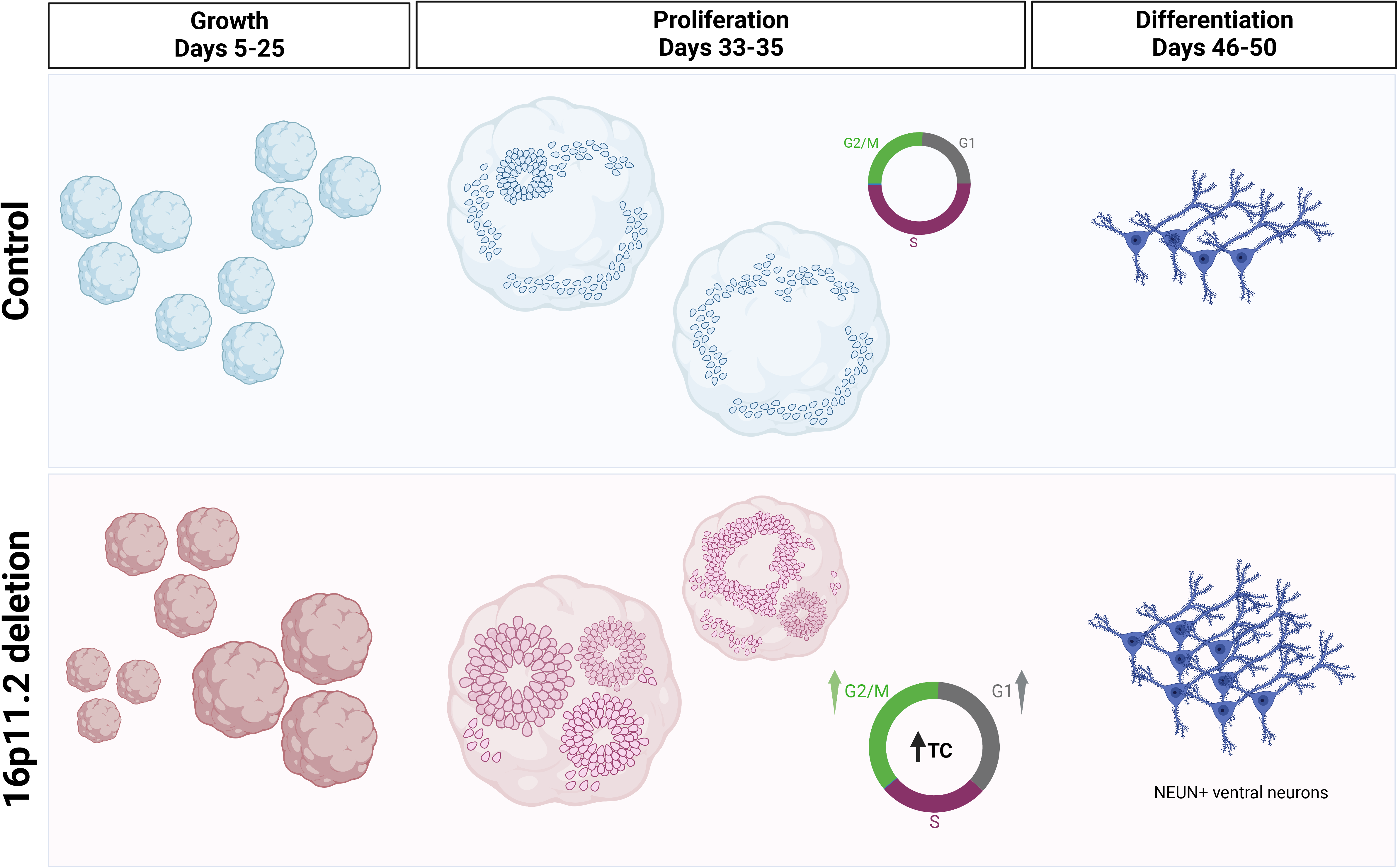
Summary of the effects of 16p11.2 deletion in ventral organoids. Control organoids from the different lines grew at similar rates, whereas deletion organoids exhibited significant variations in growth rates. Regardless of organoid size, deletion organoids from both lines displayed a greater potential to form rosettes without altering the numbers of cycling progenitors at early timepoints, with increased variability in the observed phenotype. However, cycling progenitors in the deletion organoids proliferated slower than the controls. The effect of this prolonged cell cycle length, as well as the duration of G1 and G2M, becomes evident at later timepoints where the expression of neuronal marker NEUN is increased, suggesting that this CNV accelerates neuronal production. Figure created by biorender.com.

### Variability as a hallmark of 16p11.2 deletion

In this study, deletion organoids consistently exhibited increased variability compared to their isogenic control organoids. This was evident by the increased variability observed in rosette area, COUPTFII expression and cell cycle kinetics among deletion organoids. The very similar batch-batch variation observed in both deletion lines (**Fig.S2, S3**) argued against a significant contribution from the heterozygous duplication affecting six protein coding genes on chromosome 4 in deletion line DELD5 and any other genetic differences between these effectively isogenic lines.

This variability is important when considering why 16p11.2 deletion patients also demonstrate a variable clinical phenotype. The clinical heterogeneity and incomplete penetrance in 16p11.2 patients are indeed quite remarkable, thus rendering the correct clinical diagnosis challenging (Fetit et al., 2020). We propose that the loss of this region, which contains several genes that are involved in the regulation of the cell cycle during neurogenesis and converge on cytoskeletal and cell adhesion pathways such as *MAPK3, MVP* and *KCTD13* (Golzio et al., 2012, Pucilowska et al., 2015, Roth et al., 2020), unleashes the constraints present throughout development, introducing variability and increasing the range of possible outcomes as to which developmental course the cells undertake. In humans, this divergence from neurotypical development should not, in principle, follow predictable routes, but is rather largely dependent on initial starting differences, such as the patient’s genetic background, recessive mutations in the non-deleted homologue or the variability in size of the lost fragment (Pizzo et al., 2019, Coman and Gardner, 2007, Szelest et al., 2021), which translates into the remarkable phenotypic variability observed in 16p11.2 patients. Using isogenic cell lines in our ventral organoid model of 16p11.2 deletion eliminates the variability which may arise due to differences in genetic background or due to the size of the deleted fragment. Rather, the deletion of this locus appears to introduce instability into the system, rendering the organoids more sensitive to external culture conditions, as evident by how the manifested effects varied between batches but not according to deletion cell line.

The variability in early growth rates observed in the deletion organoids in this study recapitulates the variable effects of 16p11.2 deletions on sub-cortical MGE and CGE-derived structures reported in the current literature. A recent brain MRI study revealed increased volumes of the basal ganglia structures, putamen and pallidum, in patients with the deletion of the distal 16p11.2 region (Sønderby et al., 2020), whereas another neuroimaging study of the deleted proximal 16p11.2 region reported no alterations in basal ganglia structures (Martin-Brevet et al., 2018). Reports on the incidence of macrocephaly among individuals with 16p11.2 deletion are equally variable and range from 17% to 69% (Steinman et al., 2016) (Qureshi et al., 2014) (Zufferey et al., 2012). Therefore, the variability in ventral organoid size reflects the variation in 16p11.2 and ASD patients with respect to basal ganglia size and overall brain size. The current reports on the effects of 16p11.2 deletion on cortical NPC proliferation are few and variable, leaving the effects of this CNV on the ventral telencephalon largely unexplored. Enhanced proliferation (Connacher et al., 2022, Pucilowska et al., 2015), no difference in NPC proliferation rates (Deshpande et al., 2017, Roth et al., 2020) or a reduction in NPC proliferation (Urresti et al., 2021), as observed in our ventral progenitors, have been reported, further highlighting the increased variability associated with this deletion.

### Increased rosette organisation suggests premature differentiation in 16p11.2 deletion organoids

Deletion organoids exhibited enhanced rosette formation which could be a sign of premature differentiation. NPCs progress from unstructured neuroepithelial cells to form rosettes *in vitro* (Ziv et al., 2015) and are capable of responding to different patterning cues to initiate the differentiation into region-specific neuronal fates (Elkabetz et al., 2008). The increased potential of deletion organoids to form rosette might, therefore, render the ventral progenitors more likely to respond better to differentiation signals.

Transcriptomic profiling of cortical neural progenitor cells derived from human iPSCs harbouring the deletion revealed several differentially expressed genes, both within and outside the 16p11.2 locus, involved in cytoskeletal organization and cell adhesion (Roth et al., 2020). Because rosettes are formed through cytoskeletal events (Harding et al., 2014), we can speculate that the effects of this deletion may converge on cytoskeletal pathways that promote the rearrangement and organisation of ventral progenitors into neural rosettes, without altering the number of neural progenitors within the organoid at earlier stages. Roth et al., 2020 reported no observable differences in the abilities of control and 16p11.2 deletion cortical NPCs to form rosettes, although this finding was not quantified (Roth et al., 2020). In contrast to our findings, the absence of a phenotype in cortical NPCs may suggest that 16p11.2 deletion could affect dorsal and ventral progenitors differently.

COUPTFII is another factor that is an important regulator of differentiation during embryonic development (Polvani et al., 2019) and plays a role in the formation of neural rosettes *in vitro* (Fedorova et al., 2019, Hsu et al., 2017), raising the possibility that increased COUPTFII expression might affect the generation of rosettes in the deletion organoids. Conversely, the higher abundancy of rosettes in the deletion organoids might result in increased COUPTFII expression.

### Prolonged cell cycle length and G1-phase duration drive early differentiation in 16p11.2 deletion organoids

16p11.2 deletion causes ventral progenitors to proliferate slower in ventral organoids, which increases their likelihood of undergoing neurogenic cell divisions and differentiating into interneurons. Accelerated ventral telencephalic differentiation might also occur in humans with 16p11.2 deletion, as we observed in ventral organoids. Indeed, many 16p11.2 genes, such as KIF22, ALDOA, HIRIP3, PAGR1, and MAZ were found to be expressed in progenitors and could, therefore, influence neurogenesis (Roth et al., 2020, Morson et al., 2021). To date, only one study utilised organoids to examine the effect of 16p11.2 deletion on cortical NPC proliferation at one month, revealing a decreased proliferation rate and higher NEUN+ cells (Urresti et al., 2021), which is in line with our findings in ventral organoids.

The increased cell cycle length in the day 33-35 deletion organoids could play a role in the increased NEUN expression observed in days 46 and 50 deletion organoids. Eventually, the increased withdrawal of cells from the cell cycle and the depletion of the progenitor pool might explain why no differences in NEUN levels were observed between deletion and control organoids at day 70. During the G1-phase, cells sense different environmental cues to initiate cell-fate decisions (Lanctot et al., 2017, Florio and Huttner, 2014), and its lengthening promotes the transition to a more differentiated progeny (Dalton, 2015, Hwang et al., 2021). The lengthening of G1-phase in cortical progenitors is associated with the fate-transition from apical progenitors to basal progenitors, and increases the likelihood that a daughter cell exits the cell cycle, thus promoting neurogenesis (Lim and Kaldis, 2012, Arai et al., 2011, Pilaz et al., 2009, Lange et al., 2009). Labelling experiments in the mouse LGE have demonstrated a progressive lengthening of G1 with embryonic age (Bhide, 1996, Sheth and Bhide, 1997). Moreover, recent studies provided evidence for a causal link between the cell cycle length of MGE and LGE progenitors and the cell fate of their progeny (Zong et al., 2022, Magnani et al., 2010). Thus, the altered cell cycle length and increased NEUN expression in our ventral deletion organoids is compatible with the possibility that this deletion may result in increased ventral neuron production.

## Conclusion

In conclusion, we propose the following hypothesis based on our findings: at early stages of ventral telencephalic development, the 16p11.2 deletion enhances rosette formation in ventral organoids without altering the proportions of neural progenitors or immature neurons within the organoid. Because neural progenitors organized into rosettes can respond to environmental cues and initiate differentiation into region-specific neuronal fates better than dispersed progenitors, the more-structured NPCs in the deletion organoids are more likely to undergo neurogenic cell divisions. The prolongation of total cell cycle length as well as the duration of G1-phase also increases the likelihood of ventral progenitors differentiating into neurons, which eventually becomes evident at later timepoints. Our findings also indicate that the 16p11.2 deletion mechanism introduces more variability into this developing system.

## Materials and Methods

### Culturing of iPSC lines

Cells were generated and provided by (Tai et al., 2016). Briefly, isogenic control lines were generated by transfecting the parent iPSC line, GM8, with the Cas9 expression vector lacking any guide-RNAs as to not induce any genetic modifications. All deletion lines were generated by transfecting the parent iPSC line with the Cas9 expression vector, including the designed guide RNAs to target the homologous sites flanking the 16p11.2 locus. This method generated a 740-kb microdeletion of the 16p11.2 region that mimics the consequences of the in vivo non-allelic homologous recombination (NAHR) and mirrors the size of the CNV in humans as previously described (Tai et al., 2016).

A total of 7 iPSC lines were used: The parent line, GM8 and three isogenic control lines (FACS51, FACS52 and FACS53) together with three deletion lines (DELD5, DELA3 and DELB8). Cells were cultured in feeder-free medium and grown on Matrigel-coated plates in cell medium containing 1:1 mTesR1: Essential 8 (StemCell Technologies, 85850 and Thermofisher A1517001, respectively). The cells were maintained so that once the iPSC colonies were confluent, cells were split and passaged into different wells. The cells were maintained and passaged until stable, growing into healthy colonies and showing very little differentiation before being used to grow organoids.

### Generating ventral organoids

To generate ventral organoids, we adapted the pre-established protocol developed by (Sloan et al., 2018). Briefly, once iPSC lines were confluent, differentiated cells were manually removed. Accutase (Stem Cell Technologies, #07920) was added (1ml/well) and incubated for 5-6 minutes to detach the cells from the plate. Cells were washed with PBS and resuspended in 1ml E8:MTESER media with ROCK inhibitor Y-27632 (10μM, EMD Chemicals). 9000 cells were seeded per well in a 96-well round bottom ultra-Low attachment plate (Corning). The cells remained in E8:MTESER media with ROCK inhibitor for a transition period of 4 days, the media was replenished once on the second day. This increased the chances of the cells aggregating into organoids. Neuronal induction media (NIM) and neuronal media (NM) were prepared as described in (Sloan et al., 2018) and summarised in **Table S8**. Following this transition period, individual organoids were transferred from 96-well plates to 24 well plates in NIM supplemented with two SMAD inhibitors; dorsomorphin (5μM, Sigma-Aldrich) and SB-431542 (10μM, Tocris), together with ROCK inhibitor Y-27632 (10μM, EMD Chemicals). Growing organoids in separate wells, using the same number of cells seeded per well eliminates the issue of organoid fusion and allows for a fair comparison of organoid size.

Plates were then incubated at 37 degrees and 5% CO_2_ for 48 hours. On the 3rd day, fresh NIM supplemented with dorsomorphin, SB, and the Wnt pathway inhibitor IWP-2 (5μM, Selleckchem) was added and media was changed every day. On day 6, organoids were transferred to NM containing neurobasal-A (Life Technologies, 10888), B-27 supplement without vitamin A (Life Technologies, 12587), GlutaMax (1:100, Life Technologies), penicillin and streptomycin (1:100, Life Technologies) and supplemented with the growth factors EGF (20 ng ml^−1^; R&D Systems) and FGF2 (20 ng ml^−1^; R&D Systems) until day 24. From day 6 onwards, the cells were also incubated on an orbital shaker.

To induce ventral identity, SHH pathway agonist SAG (smoothened agonist, 100 nM, Selleckchem) was added to NM from day 12 to day 24. IWP-2 (Selleckchem#s7085) and SAG as well as allopregnanolone (100 nM, Cayman Chemicals) were supplemented from day 15 to day 24 with a brief exposure (day 12–15) to retinoic acid (100 nM, Sigma-Aldrich). From day 25 to 42, NM was supplemented with the growth factors BDNF (20 ng ml^−1^, Peprotech) and NT3 (20ng ml^−1^, Peprotech) and media was changed every other day. From day 43 onwards, organoids were maintained in unsupplemented NM with medium changes every four to six days.

### Cryopreservation and Immunohistochemistry

Organoids were fixed in 4% paraformaldehyde (PFA) for 30 min to 2h. They were then washed in PBS 3 times and transferred to 30% sucrose solution overnight at 4 °C. After that, they were transferred into embedding medium containing 1:1 30% Sucrose:OCT, snap-frozen on dry ice and stored at −80 °C. For IHC, 10-μm-thick sections were cut using a cryostat (Leica). Cryosections were warmed at room temperature and washed in running water, followed by washing in 0.1% Triton X-100 diluted in PBS (PBST) to permeabilize the tissue. Sections were then blocked in 10% donkey or goat serum in PBST for 30 minutes at room temperature. After that, they were incubated overnight at 4 °C with primary antibodies diluted in blocking solution, as listed in **Table S8**. PBST was used to wash off the primary antibodies and the cryosections were incubated with secondary antibodies in blocking solution for 1 h. Finally, nuclei were visualized with DAPI and Cryosections were mounted for microscopy on glass slides using Vectashield Hardset (Vector Labs) for imaging on a fluorescent or confocal microscope. Images were processed in ImageJ (Fiji).

### Measuring Organoid Size

Organoids were imaged in culture using a light microscope at several timepoints across development. Images were then analysed using ImageJ (Fiji). The organoid outline was selected using the polygon selection tool and the area was measured for every organoid image. The individual organoid area was then normalised to the average area of the control organoids in the respective batch. Boxplots and line graphs were plotted using R.

### Quantification of neural rosettes

Images were taken at 10x to cover the entire organoid and analysed using ImageJ (Fiji). The organoid outline was selected using the polygon selection tool and the area was measured for every organoid image. Similarly, neural rosettes were outlined, and their total area was measured. The number of neural rosettes per organoid was quantified, together with the total area occupied by the neural rosettes in an organoid. The relative rosette area was also calculated by dividing the total area occupied by neural rosettes in an organoid by the area of the organoid.

### Quantification of COUPTFII expression

Images were taken at 10x to cover the entire organoid and analysed using ImageJ/Fiji. The images were then split into the different channels to quantify the corresponding markers. Cells positive for COUPTFII were manually quantified using the manual cell counter plug-in in ImageJ/Fiji.

### Dissociation of organoids for flow cytometry

Organoids were transferred to 6-well plates and washed twice with PBS. Next, the organoids were incubated with 2ml Accumax (Stem Cell Technologies, #07921) for 10 minutes at 37 degrees on an orbital shaker (90rpm). Using a p1000 tip, organoids were manually dissociated by pipetting up and down then centrifuged at 200g for 5 minutes. The pellet was then resuspended in 1ml PBS and filtered through a 40um filter into 1.5ml Eppendorf tubes. Cell density and viability were quantified.

### Fixation and permeabilization of cells

FOXP3 Fix/Perm Buffer set (Biolegend, #421403) was used according to manufacturer’s instructions. Briefly, 1ml of 1X Biolegend’s FOXP3 Fix/Perm solution was added to each tube, vortexed, and incubated at room temperature for 20 minutes. Samples were centrifuged, the supernatant removed then washed once with FACS staining buffer (ThermoFisher, #00-4222-24). Cells were then resuspended in 1ml 1X Biolegend’s FOXP3 perm buffer and incubated in the dark at room temperature for 15 minutes. After that, the samples were centrifuged and resuspended in 100ul of 1X FOXP3 Perm buffer. A master mix of the primary conjugated antibodies for SOX2, Ki67 and TUJ1 together with Hoechst 33342 was prepared in FACS staining buffer (**Table S8**) and the cells were incubated with the master mix for 1 hour in darkness at room temperature. Single staining controls were prepared for every antibody used as well as an unstained control. Cells were washed once with FACS staining buffer and resuspended in 300-500ul FACS staining buffer for analysis. with the flow cytometer. Samples were analysed using the BD LSRFortessa™ cell analyser at the QMRI flow cytometry and cell sorting facility, University of Edinburgh.

### Flow cytometric Analysis

Data was processed using FlowJo v10.7.2. Briefly, single-stained and unstained controls were used to identify and set thresholds for our gates. A combination of density plots and contour maps for forward scatter (FSC) was used to clearly outline the cell populations positive for the individual markers, particularly where no clear single histogram peaks could be identified. Doublets were excluded, first, by plotting Hoechst-Area versus Hoechst-Height to eliminate G0 and G1 doublets from the rest of the cells in the cell cycle and using the FSC-Area versus FSC-Height to further confirm that only single cells were included. From the parent single cell population, TUJ1+ neuron and SOX2+ progenitor populations were then identified and quantified. The proportion of SOX2+Ki67+ cells (proliferative pool) were also quantified from the population of SOX2+ cells. From the SOX2+ progenitors, cells positive for Ki67 were plotted against Hoechst 33342 to differentiate the cells based on their DNA content and quantify the number of cells in the different cell cycle phases (Fig. S6). Finally, results were exported to be processed in R.

### IdU/BrdU double labelling

IdU/BrdU double labelling was done as described by (Martynoga et al., 2005) by sequentially exposed our organoids to the halogenated thymidine analogues: iododeoxyuridine (IdU) and bromodeoxyuridine (BrdU), which are incorporated into the synthesized DNA during S-phase. Briefly, IdU and BrdU labelling solutions were freshly prepared in IdU/BrdU solvent (0.9% NaCl). Organoids were transferred to a 5cm dish. A first pulse of 100uM IdU labelling solution was added to 5 ml of the media and incubated for 1.5 hours at at 37 degrees and 5% CO2 on an orbital shaker (45rpm). This labels all the progenitors in S-Phase (S_cells_) at the beginning of our experiment. After 1.5 hours, the organoids are then exposed to a pulse of 100uM of the BrdU labelling solution and incubated on an orbital shaker for 30 minutes to make sure that all BrdU was well-incorporated into the progenitors then fixed for analysis. This labels S_cells_ at the end of the experiment. Because neural progenitors are not synchronized as they go through the cell cycle (Takahashi et al., 1993), the initial cohort of IdU-labelled cells will exit the S-phase at a constant rate and will constitute the fraction of leaving cells (Lcells) that are only positive for IdU but not BrdU. In this experimental set-up, the time interval during which the cycling progenitors can incorporate only IdU (Ti) is 1.5 hours. Finally, organoids were fixed for staining and processed for IHC analysis as described above. Using antibodies that allow for the differentiation between cells labelled with IdU only and those labelled with both IdU and BrdU (**Table 1**), we quantified the Lcells and S_cells_.

### Quantification of cell cycle kinetics

Calculation of the total cell cycle length (TC) and the duration of G1 (T_G1_), S (Ts) and G2M (T_G2M_) phases was done as described by (Martynoga et al., 2005) and (Mi et al., 2013). Briefly, because it has been shown that the ratio of the length of an individual phase of the cell cycle to that of another phase equals the ratio of the cell numbers in the two phases (Nowakowski et al., 1989), the ratio between T_i_ (1.5 hours) and T_s_ (duration of the S-phase) equals the ratio between L_cells_ (IdU+BrdU−) and S_cells_ (IdU+BrdU+) as shown below:

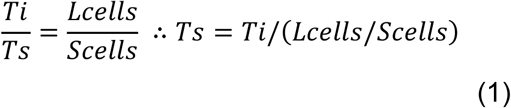

The fractions of L_cells_ (IdU+BrdU−) and S_cells_ (IdU+BrdU+) were first quantified and Ts was calculated as per *Formula (1)*. **Table S5** lists the numbers of L_cells_ and S_cells_ calculated from the double IdU/BrdU labelling experiment for the organoids analysed in the different cell lines across the 4 batches. The number of proliferating progenitor cells, which we refer to as the proliferative pool (SOX2+Ki67+) and calculated from flow cytometry, was denoted as P_cells_. **Table S6** summarises the average numbers of P_cells_, together with the average number of cells in G1, S and G2M phases calculated from all organoids analysed using flow cytometry.

The total cell cycle length (T_c_) can then be calculated using the ratio between T_s_ and T_c_ which equals the ratio between S_cells_ and P_cells_, as shown below:

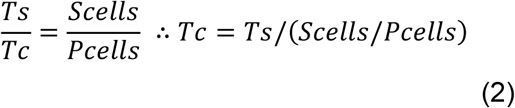

Having already calculated T_s_ from the double labelling experiment using *Formula(1)*, we used the values of S_cells_ and P_cells_ calculated from the flow cytometry (**Table S6**) to calculate TC using *Formula (2)*. Once T_c_ was calculated, the duration of cells in G1 (T_G1_) was calculated as per *Formula (3)*:

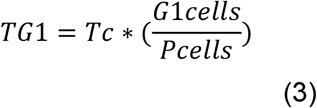

Where G1_cells_ is the number of cells in G1 phase and P_cells_ is the number of cycling cells in our progenitor pool. Dividing G1_cells_ by P _cells_ gives us the fraction of cells in G1 phase. Multiplying this value by total cell cycle length provides us with the duration of G1 (T_G1_). Finally, having calculated Tc, Ts and T_G1_, we were able to calculate T_G2M_ by simple subtraction, as shown in Formula (4):

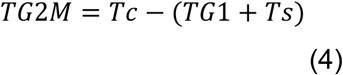

### RNA extraction

Organoids were loaded onto the QIAshredder (Qiagen, #79654) homogenizer to homogenize the tissue and the lysate is then collected. RNA was then extracted using RNeasy Mini Kit (Qiagen, #74104) according to manufacturer’s instructions. RNA waseluted in a total of 20ul RNase free water. The extracted RNA was quantified using Nanodrop and prepared for real time quantitative polymerase chain reaction (RT-qPCR).

### Real-time quantitative PCR (RT-qPCR)

cDNA was prepared using LunaScript^®^ RT SuperMix Kit (New England Biolabs, #E3010L), together with no-RT control reaction and no template controls, according to manufacturer’s instructions. Reactions were then incubated in the thermocycler at 25° for 2 minutes for primer annealing, 55° for 10 minutes for cDNA synthesis and finally at 95° for 1 minute for heat inactivation. cDNA was then diluted to a final concentration of 1ng/μl or 1ng/10 μl reaction for samples with lower concentrations. RT-qPCR was performed using Luna® Universal qPCR Master Mix. Primers sequences are listed in **Table S8** and adapted from (Birey et al., 2017). ΔΔCt method was used to normalise and quantify relative fold changes in gene expression to two housekeeping genes.

### Quantification of NEUN intensity

Images were taken at 5x to cover the entire organoid and analysed using ImageJ/Fiji. The images were then split into the different channels to quantify the corresponding markers. The polygon tool was used to delineate the organoid and measure its area. The mean grey value was measured to represent the mean intensity of the NEUN+ cells within the organoid. The relative NEUN intensity was calculated by dividing the mean NEUN intensity by the organoid area.

### Data Analysis and Statistics

Analysis was performed using R studio (versions 3.3.2 and 4.0.4). Graphs were plotted using the R package (ggplot2). Levene’s Test in the R package (car) was used to assess the homogeneity of variance. Because our data did not fulfil the homoscedasticity assumption, the non-parametric Welch’s ANOVA was used to assess statistical significance. Post-hoc comparisons were performed using Games Howell test, an alternative to Tukey’s comparisons, in the R package (rstatix). Finally, the effect size was calculated using the non-parametric Cohen’s d function in the R package (effsize).

Statistical analysis was performed on the flow cytometric dataset using Linear Mixed Effects (LME) modelling to account for the variation introduced into our findings due to batch effects. This was followed by Type-III ANOVAs to attain p-values and assess the statistical significance of LME models. The models generated were both random intercept models, to account for baseline-differences in batches, and random slope models, to accommodate that the effect of different cell lines might be different for different batches. The R package (lmerTest) was used to design and generate LME models. The R package (car) was then used to run ANOVA tests on our models to assess the statistical significance of our findings. Finally, the R package (sjstats) was used to calculate the effect sizes using omega_squared function.

## Acknowledgements

We would like to thank James Gusella and Derek Tai at the Molecular Neurogenetics Unit, Harvard for providing the cell lines used in this study, Michela Barbato and Wai-Kit Chan (Calvin) for their help and advice on maintaining the cell lines and generating organoids and on imaging, Yu-Ting Huang (Nikky) for her input on designing the flow cytometry experiment and, finally, Shonna Johnston and Will Ramsay at the QMRI flow cytometry and cell sorting facility, University of Edinburgh for their guidance and support whilst processing the samples and analysing the flow cytometry data, Nicola Wrobel and Lee Murphy at the Edinburgh Clinical Research Facility, Western General Hospital for running the CytoSNP array.

